# Identification and characterization of *DICER-LIKE* genes and their roles in *Marchantia polymorpha* development and stress adaptation

**DOI:** 10.1101/2023.02.03.526932

**Authors:** Erika Csicsely, Anja Oberender, Anastasia-Styliani Georgiadou, Nora Gutsche, Sabine Zachgo, Oguz Top, Wolfgang Frank

## Abstract

DICER-LIKE (DCL) proteins have a central role in plant small RNA (sRNA) biogenesis. The *Marchantia polymorpha* genome encodes four DCL proteins: two DCL1 homologs, MpDCL1a and MpDCL1b, MpDCL3 and MpDCL4. While MpDCL1a, MpDCL3 and MpDCL4 show high similarities to their orthologs in *Physcomitrium patens* and *Arabidopsis thaliana*, MpDCL1b shares only a limited homology with PpDCL1b, but it is very similar, in terms of functional domains, to orthologs in *Anthoceros agrestis* and *Salvinia cucullata*. We generated Mp*dcl*^*ge*^ mutant lines via the CRISPR/Cas9 system and performed comprehensive phenotypic analyses of these mutant lines, under control and salt stress conditions as well as upon exogenous naphthaleneacetic acid (NAA) and abscisic acid (ABA) treatments to gain insights into the respective MpDCL functions. While Mp*dcl1a*^*ge*^ mutants display severe developmental aberrations throughout their development, no adverse phenotypic changes are detectable in Mp*dcl1b*^*ge*^ and Mp*dcl4*^*ge*^ mutants except the development of less and smaller male sexual organs (antheridiophores) when they are cultivated under photoperiod conditions supplemented with far-red light. Mp*dcl3*^*ge*^ mutants display rosette-shaped thallus formation and overall faster development, but are not able to form antheridiophores. The rosette-shaped thallus development of Mp*dcl3*^*ge*^ can be reverted to a wild-type-like thallus growth upon NAA treatment. Mp*dcl1b*^*ge*^ mutants can tolerate high levels of salt, whereas Mp*dcl4*^*ge*^ mutants show higher salt sensitivity. Moreover, Mp*dcl1a*^*ge*^ and Mp*dcl3*^*ge*^ mutants show an ABA-hypersensitive phenotype. It can be concluded that the observed phenotypic alterations, under normal or treatment conditions, are linked to the mutations in the respective Mp*DCLs* and hence to defective or altered sRNA biogenesis pathways in *M. polymorpha*. In conclusion, MpDCLs and their associated sRNAs regulate development, abiotic stress and phytohormonal response in *M. polymorpha*.

## Introduction

The ancestors of today’s modern bryophytes were among the first to conquer land as a habitat (Degola *et al*., 2022). When these early land plants transitioned to life on land some 450–470 million years ago, they acquired new innovations to survive in their new habitat. These include a rapid extension of the regulatory function of small non-coding RNAs (sRNA) that recognize reverse complementary RNA or DNA sequences to mediate posttranscriptional and transcriptional gene silencing (Fattash *et al*., 2007, Arif *et al*., 2013). Further, different classes of sRNAs, microRNAs (miRNA) and different classes of small interfering RNA (siRNA), evolved that control multiple diversifications of plant body plans, developmental mechanisms and adaptations to new abiotic stress factors (Fattash *et al*., 2007). As a result, proteins that act in sRNA biogenesis emerged through parallel evolution of the small RNA machinery in different Chlorophyta lineages, including DICER-LIKE proteins (DCL) (Margis *et al*., 2006, Fattash *et al*., 2007, Cui *et al*., 2017, Bélanger *et al*., 2022, Bélanger *et al*., 2023).

The most common plant DCL1 protein evolved from an algae-typical DCL after the divergence of Klebsormidiophyceae and other Streptophyta lineages (Wang *et al*., 2021). DCL3 and 4 emerged in the bryophyte lineages (You *et al*., 2017, Wang *et al*., 2021, Bélanger *et al*., 2022, Bélanger *et al*., 2023) while DCL2, another common DCL in embryophytes, originated in the last common ancestor of ferns and seed plants (You *et al*., 2017, Bélanger *et al*., 2022, Bélanger *et al*., 2023). Additionally, most monocot plant lineages like rice and maize express *DCL5* homologs (Patel *et al*., 2021, Bélanger *et al*., 2022, Bélanger *et al*., 2023). Proteins of the plant DCL family are involved in the biogenesis of different sRNA classes like miRNAs, trans-acting small interfering RNAs (ta-siRNAs), heterochromatic small interfering RNAs (hc-RNAs), phased small interfering RNAs (phasiRNAs), repeat-associated small interfering RNAs (ra-siRNAs), and natural antisense transcript-derived small interfering RNAs (nat-siRNAs) (Arif *et al*., 2013, Habermann *et al*., 2020, Tiwari *et al*., 2021).

The *DCL* gene family usually contains numerous members that vary among plant species, with four to five members being most common among seed plants and at least one *DCL* gene in most green algae species (Margis *et al*., 2006, Liu *et al*., 2009, Bélanger *et al*., 2022, Bélanger *et al*., 2023). Although the core mechanisms of DCL function are evolutionarily conserved, multiple pathways with shared components and overlapping functions that generate sRNAs of specific sizes with dedicated functions have evolved especially in seed plants (Liu *et al*., 2009, You *et al*., 2017, Yu *et al*., 2017, Tiwari *et al*., 2021). One *DCL* gene found in all land plant species encodes DCL1 whose function in miRNA biogenesis is highly conserved (Arif *et al*., 2013, Yu *et al*., 2017, Manavella *et al*., 2019, Arif *et al*., 2022). During plant miRNA biogenesis, *MIR* genes are transcribed by RNA polymerase II and the primary miRNA (pri-miRNA) folds back into a hairpin-like structure that is recognized by a multiprotein microprocessor complex including DCL1, RNA double-strand binding protein Hyponastic Leaves 1 (HYL1) and the zinc finger protein SERRATE 1 (SER1). After processing of the pri-miRNA into a precursor miRNA (pre-miRNA), DCL1 and HYL1 produce a specific miRNA/miRNA* (miRNA strand/passenger strand) duplex from the foldback region. This duplex is 2’-O-methylated at the 3’ ends by Hua Enhancer 1 methylase (HEN1) to protect it from degradation after transport into the cytoplasm (Baranauske *et al*., 2015, Pietrykowska *et al*., 2022). The miRNA guide strand is recruited into an Argonaute 1 (AGO1) protein that is part of the RNA-induced silencing complex (RISC), and exported into the cytoplasm with the help of Chromosomal Maintenance 1 (CRM1) (Arif *et al*., 2013, Yu *et al*., 2017, Bologna *et al*., 2018, Zhang *et al*., 2020, Tiwari *et al*., 2021). The incorporated miRNA guides the RISC to complementary targets which results in posttranscriptional gene silencing (PTGS) via RNA cleavage and translational inhibition or transcriptional gene silencing (TGS) via DNA and/or histone modifications. Plant DCL proteins are characterized by DEAD-box, Helicase-C, DUF283, PAZ (Piwi/Argonaute/Zwille), Ribonuclease 3 (RNase III), and double-stranded RNA binding (dsRBD) domains (Margis *et al*., 2006, Liu *et al*., 2009), but not each plant DCL harbors all domains. Most plant DCLs have a different set of these domains that might indicate interchangeable and/or specific functions that underlie the origin and evolution of DCLs in land plants (Margis *et al*., 2006). It is proposed that a functional DCL protein should contain an RNA binding site, a PAZ domain as well as two RNase III (RNase III a and b) domains found in close proximity to each other forming the active center of the protein (Margis *et al*., 2006, Liu *et al*., 2009, Liu *et al*., 2012).

sRNAs are not only known to regulate and initiate responses to abiotic and biotic stresses (Willmann and Poethig, 2007, Barciszewska-Pacak *et al*., 2015, Tiwari *et al*., 2020, Tiwari *et al*., 2021), but they also control plant development and organ formation (Willmann and Poethig, 2007). For instance, miR160, miR164, miR167, miR390, and miR393 control the expression of auxin response factor genes (*ARF*) that encode transcription factors which participate in *Arabidopsis thaliana* root development and stress response (Meng *et al*., 2010). In addition, *A. thaliana* miRNAs are differentially expressed in response to various abiotic stresses such as dehydration, salt and ABA (Sunkar and Zhu, 2004), high light (Tiwari *et al*., 2021) and cold (Tiwari *et al*., 2020) and were also shown to act in retrograde signaling (Habermann *et al*., 2020). Furthermore, miR160 targets Mp*ARF3* in the bryophyte *Marchantia polymorpha* and Mp*miR160* overexpression results in major developmental aberrations including auxin insensitivity, constitutive tropisms and misregulated cell division patterns during gemmae development comparable to the phenotype of the loss of function Mp*arf3* mutant line (Flores-Sandoval *et al*., 2018a). Similarly, miR160, a miRNA that is conversed across land plants, regulates other *ARF* genes in many plants like *OsARF18* in rice, *AtARF10, AtARF16* and *AtARF17* in *A. thaliana* and *PpARF* in *Physcomitrium patens* (Arazi, 2012, Huang *et al*., 2016, Jodder, 2020).

While sRNA biogenesis and the involvement of various DCL proteins in this process are well examined in seed plants like *A. thaliana*, the functions of DCLs in the liverwort *M. polymorpha* have not been elucidated so far (Pietrykowska *et al*., 2022). Like *A. thaliana*, the bryophytes *P. patens* and *M. polymorpha* encode four DCL proteins. They both encode two DCL1 homologs (DCL1a and DCL1b), DCL3, and DCL4 (Arif *et al*., 2013, Lin and Bowman, 2018) whereas a DCL2 homolog is missing. In previous studies, we uncovered the specific functions of all four *P. patens* DCL proteins in the biogenesis of different sRNA classes and their impacts on various developmental processes (Cho *et al*., 2008, Khraiwesh *et al*., 2010, Arif *et al*., 2012). We also recently elucidated the involvement of an miR1047-based feedback control of *PpDCL1a* transcript abundance which maintains a balance between intronic miRNA biogenesis and proper *PpDCL1a* mRNA splicing. Furthermore, the miR1047-based autoregulatory feedback loop is essential for salt tolerance and its abolishment results in global changes in the mRNA and miRNA expression (Arif *et al*., 2022). It was proposed in a previous study that DCL1 emerged in the last common ancestor of Klebsormidiophyceae and Streptophyta algae while DCL2/3/4 diverged during the evolution of embryophytes (You *et al*., 2017, Wang *et al*., 2021). Hence, during the course of evolution and diversification, functional specialization of members of sRNA biogenesis pathways is expected. Despite homologies, MpDCLs might have some unique functions other than their counterparts in other plant species including *P. patens* and *A. thaliana*.

To gain further insight into the evolution of DCL proteins and their role in sRNA biogenesis in *M. polymorpha* we generated mutant lines for all four Mp*DCLs* and analyzed developmental abnormalities under control growth conditions, and upon exogenous naphthaleneacetic acid (NAA), abscisic acid, (ABA) and NaCl treatments. These comprehensive phenotypic analyses revealed that mutations in Mp*DCL1a* and Mp*DCL3* lead to ABA hypersensitivity and severe developmental defects throughout their development. On the other hand, mutations in Mp*DCL1b* and Mp*DCL4* resulted in developmental stage specific peculiarities, especially disturbed antheridiophore formation. Surprisingly, Mp*DCL1b*^*ge*^ mutants showed increased salt tolerance. The phenotypic analyses suggest that MpDCL1a, MpDCL3 and MpDCL4 have distinct functions in *M. polymorpha* development, similar to orthologs in *P. patens* and *A. thaliana* whereas MpDCL1b, which does not share high homology with PpDCL1b and most likely did not evolve from the same last common ancestor, functions as a negative regulator of salt stress tolerance and assists antheridiophore development. Further investigations are required to elucidate the precise functions of MpDCLs and small RNA classes, that associate with certain MpDCLs, in physiological and developmental processes and how these adaptive mechanisms have contributed to plant terrestrialization.

## Results

### *In silico* analysis of *M. polymorpha* DCL proteins reveal differences in their domain architecture

All four MpDCLs were initially annotated by Lin and Bowman (Lin *et al*., 2016, Lin and Bowman, 2018) but were re-annotated by Belanger et al. (2023) who used recently released databases for their comparative analyses of DCLs from different plant species (Bélanger *et al*., 2022, Bélanger *et al*., 2023). We extended these analyses by the addition of putative DCLs from new species. We used the four *M. polymorpha* DCLs, MpDCL1a (Mp7g12090), MpDCL1b (Mp6g09830), MpDCL3 (Mp1g02840) and MpDCL4 (Mp7g11720) that all encode the essential DCL domains as queries to search for DCL homologs in established and newly emerging model organisms. By this, we identified all known, well-studied DCLs as well as putative DCLs in newly emerging model organisms: *A. thaliana* (4), *Medicago truncatula* (6), *Oryza sativa japonica* (5) and *Zea mays* (5) as well as DCL homologs in the lycophyte *Selaginella moellendorfii* (4), in two monilophyte species *Azolla filiculoides* (3) and *Salvinia cucullata* (5), the bryophytes *P. patens* (4) and *Anthoceros agrestis* (3), the streptophyte algae species *Chara braunii* (3), *Klebsormidium nitens* (1) and *Spirogloea muscicola* (3) and a chlorophyte algae species *Chlamydomonas reinhardtii* (3) (Figure 1, Supplementary Table 1). Some of these newly identified DCLs are putative homologs identified using limited publicly available bioinformatic resources and require further validation. All obtained protein sequences were used to generate a phylogenetic tree using the CLC Main Workbench (v20) applying the neighbor-joining tree method to create an unrooted circular cladogram. With this method we were able to confirm that *M. polymorpha* does not encode a DCL2 homolog, but harbors two related DCL1 proteins (MpDCL1a and MpDCL1b) similar to *P. patens, A. agrestis* and *A. filiculoides*. Based on sequence similarities we confirmed the renaming of Mp7g12090 into Mp*DCL1a*, Mp6g09830 into Mp*DCL1b*, Mp1g02840 into Mp*DCL3* and Mp7g11720 into Mp*DCL4* as proposed by Bèlanger et al. (2023) and Pietrykowska et al. (2022). All *C. braunii* and *C. reinhardtii* DCL homologs grouped together in one clade encoding specific algal-type DCL proteins (Wang *et al*., 2021). We also found that *C. reinhardtii* and *K. nitens* harbor DCLs with a reduced set of functional domains. Domain analysis via Pfam search (Pfam-A v33.1) with the *C. reinhardtii* DCL revealed the presence of two RNase III, one Dicer-dimer and one Helicase C domain while the DCL homolog of *K. nitens* encodes the additional PAZ, RES III and Dead end protein 1 homologous to double-strand RNA binding (DND1_DSRM) domain that can also be found in bryophyte DCLs (Supplementary Table 1). These structural differences support the origin of embryophytic DCL differentiation as suggested by Wang et al. (2021). On the other hand, *C. braunii* DCL G24199, unlike the hypothesis of embryophytic DCL differentiation, harbors two adjacent RNase III domains, while G34091 encodes a Dicer-dimer domain and PAZ domain. Moreover, G34093 lacks RNase III domains, but contains two RNase H-like domains and a Helicase C domain (Supplementary Table 1). According to the biochemical properties of DCL proteins and necessary domains involved in sRNA biogenesis, additional copies of *DCLs* may exist in *C. braunii* that escaped our detection using current databases. Our phylogenetic analysis also showed that bryophyte DCL3 and DCL4 do not cluster together with DCL3 and DCL4 of seed plants, respectively.

**Figure 1.**
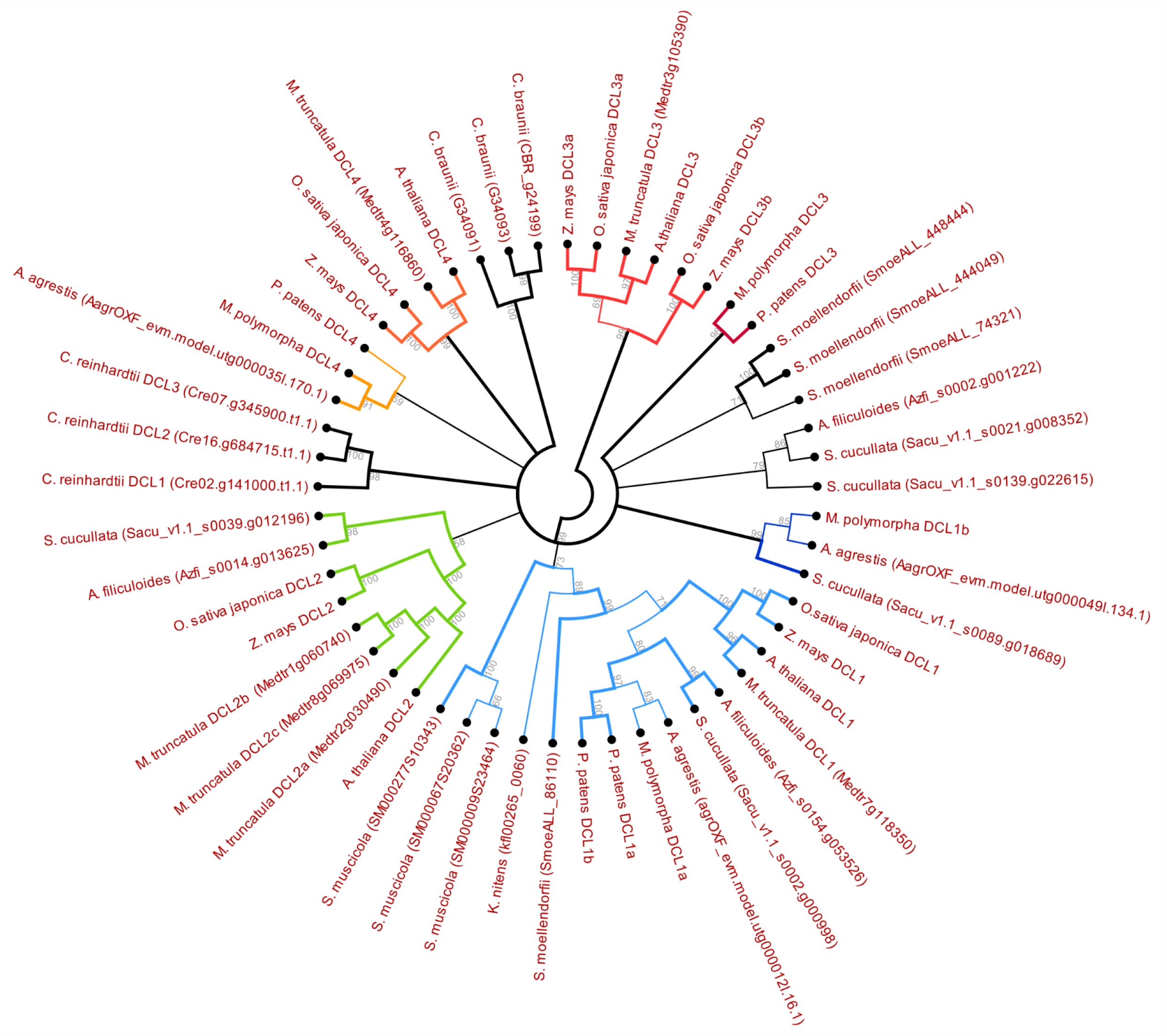
Phylogenetic analysis of DCL proteins. **(A)** Cladogram of MpDCL homologs from the following species: *Anthoceros agrestis, Arabidopsis thaliana, Azolla filiculoides, Chara braunii, Chlamydomonas reinhardtii, Klebsormidium nitens, Medicago truncatula, Oryza sativa japonica, Physcomitrium patens, Salvinia cucullata, Selaginella moellendorfii, Spirogloea muscicola*, and *Zea mays*. The accession number of the sequences is given when sequences have not been annotated previously. The cladogram was generated using CLC Genomics Workbench (Qiagen) by neighbor end joining using 1000 bootstrap replicates. Only nodes with a bootstrap value >50 are depicted and bootstrap values >90 are highlighted. Common plant DCL variants are color-coded: DCL1 homologs light blue, MpDCL1b homologs dark blue, DCL2 homologs green, bryophyte DCL4 homologs dark yellow, seed plant DCL4 homologs orange, seed plant DCL3 homologs red, bryophyte DCL3 homologs dark red, algae specific DCLs and unannotated clades black.

Detailed domain analysis of all four MpDCLs revealed that Mp*DCL1a*, Mp*DCL3* and Mp*DCL4* encode domains of RESIII, Helicase C, Dicer-dimer, PAZ and two adjacent RNase III domains. In addition to these five different domains, MpDCL1a and MpDCL4 also harbor the DND1_DSRM domain, hence they contain all six important domains (Margis *et al*., 2006, Liu *et al*., 2009, Liu *et al*., 2012). On the contrary, MpDCL1b contains only the PAZ domain and two adjacent RNase III domains (Supplementary Table 1, Figure 2A). The lack of other functional domains raises the question whether or not Mp*DCL1b* is a pseudogene. Based on our phylogenetic analysis, we found DCLs that lack some of the aforementioned six different domains. More importantly, we found potential DCL homologs with a reduced number of functional domains in *A. agrestis* (AagrOXF_evm.model. utg000049l.134.1) and in *S. cucullata* (Sacu_v1.1_s0089.g018689) which clustered together with MpDCL1b (Figure 1, Supplementary Table 1). Consequently, due to the presence of similar DCL proteins in early land plants, we hypothesize that Mp*DCL1b* may not be a pseudogene, but might have a different role than its paralogs in sRNA biogenesis.

**Figure 2.**
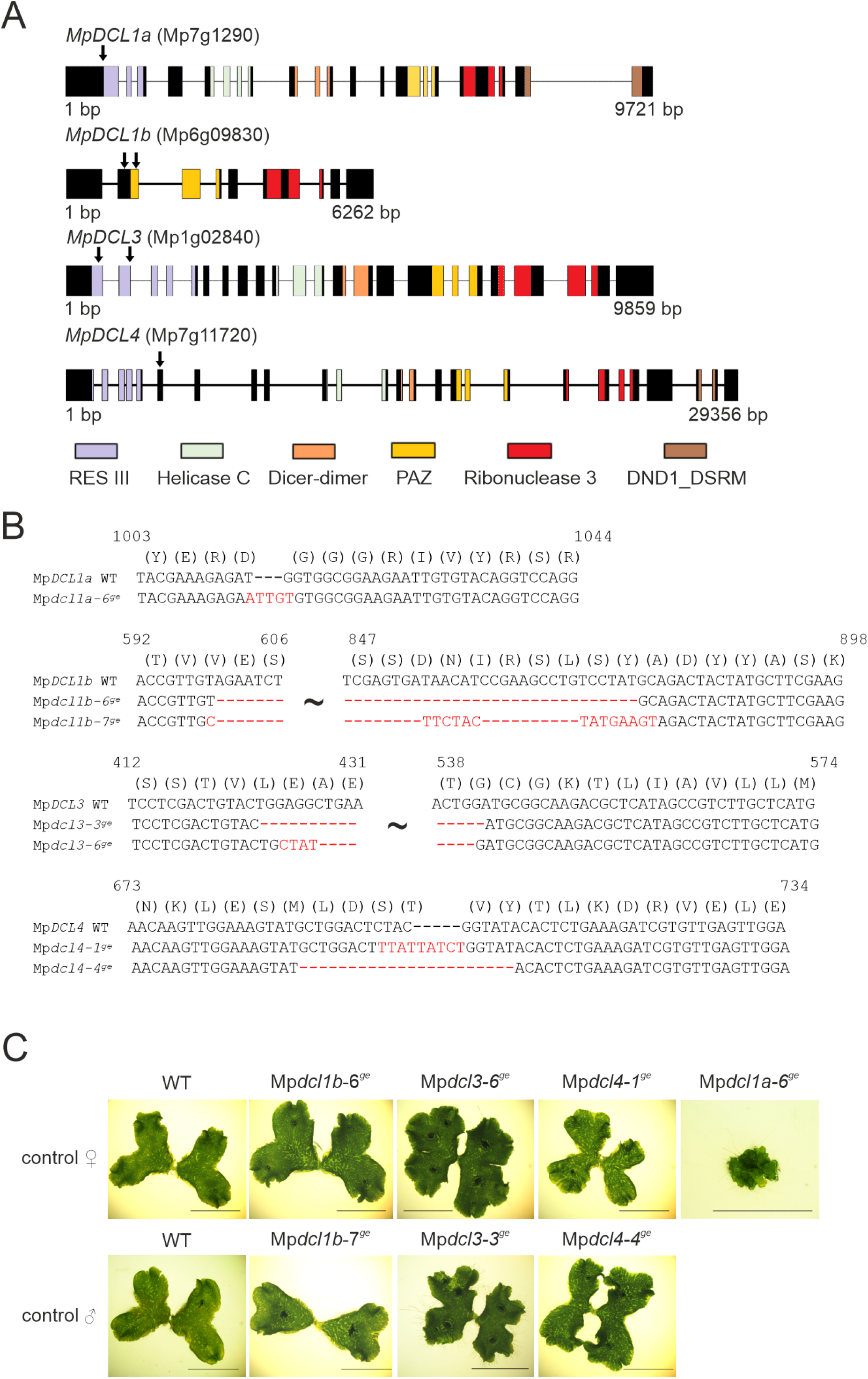
Generation of Mp*dcl*^*ge*^ mutant lines. **(A)** Schematic representation of all Mp*DCL* genomic sequences without 3’ and 5’ UTRs. The colored sections indicate encoded domains characteristic for DCL proteins. Arrows mark the targeting sites of gRNAs successfully used to generate the respective Mp*dcl*^*ge*^ mutant line. **(B)** Alignment of WT Mp*DCL* coding sequences and mutated coding sequences of ♀ Mp*dcl1a*^*ge*^ mutant line as well as ♀ and ♂ mutant lines of Mp*dcl1b*^*ge*^, Mp*dcl3*^*ge*^ and Mp*dcl4*^*ge*^; only mutated regions are shown, mutations are highlighted in red. Deletions are indicated by dashes. **(C)** Phenotypic comparison of ♀ and ♂ WT with mutant lines Mp*dcl1b*^*ge*^, Mp*dcl3*^*ge*^, Mp*dcl4*^*ge*^, and Mp*dcl1a-6*^*ge*^, 21 days after gemmae germination (DAG) and grown on standard medium under control conditions; scale bar 1 cm.

### Effects of Mp*DCL* mutations on thallus development

To investigate the functions of all four Mp*DCLs*, we generated loss-of-function mutants using the CRISPR/Cas9 system described by Sugano et al. (2018). We designed multiple gRNAs to target either the first exon or regions encoding functionally important domains of the respective Mp*DCL* gene and used these gRNAs in pairs during transformation to ensure large deletions and/or frameshift mutations. Only one gRNA from each pair induced double-strand breaks in Mp*DCL1a* and Mp*DCL4*. As a result, we obtained one male and one female line with indels that introduced premature stop codons in Mp*DCL1b*, Mp*DCL3*, and Mp*DCL4* and prevent the generation of functional proteins. The position of induced double-strand breaks within these genes as well as mutated regions are shown in Figure 2A and 2B, respectively. The use of various gRNAs in pairs (two gRNAs in each transformation event) targeting Mp*DCL1a* failed to cause large genomic fragment deletions, frameshift mutations or the generation of a premature stop codon. Analysis of all viable mutant lines displayed that only gRNA2 which did not target a genomic region encoding functional DCL domains induced a double-strand break resulting in short deletions and, in some cases, insertions of short sequences, but none of the plants surviving the selection obtained a frameshift mutation adversely affecting the encoded amino acid (aa) sequence (Figure 2A, 2B). It has been previously shown that DCL1 is an essential protein for miRNA biogenesis and is required for normal plant development. The *Atdcl1* null mutants are embryo lethal (Henderson *et al*., 2006) and the Δ*PpDCL1a* mutant lines display severe developmental disorders with altered cell size, shape and arrested development at the filamentous protonema stage (Khraiwesh *et al*., 2010). Hence, failure in generating a Mp*DCL1a* mutant line with large genomic arrangements might be due to the fact that Mp*DCL1a*, like its homologs in *P. patens* and *A. thaliana*, is essential for plant development and transformants harboring a non-functional MpDCL1a protein may not be viable. Additionally, most of the designed gRNAs were targeted to regions encoding functional DCL domains. The observed phenotypic aberrations in Mp*dcl1a*^*ge*^ mutant lines correlate with the extent of the genomic alterations. The mutant line Mp*dcl1a-6*^*ge*^ displayed the most severe phenotypic abnormalities and sequencing of the mutated region revealed the insertion of “TTG” at position 1015 and substitution of T to A (position 1014) and G to T (position 1015) (Figure 2B). These changes result in an exchange of aspartic acid (D) by glutamic acid (E) at position 338 and the insertion of leucine and cysteine (LC) at the positions 339–340 in the primary amino acid sequence of MpDCL1a. Based on the detailed analysis of the expected MpDCL1a amino acid sequence using Phyre^2^ (Kelley *et al*., 2015), we detected a potential change in the secondary structure of the protein (position 280–390) due to the aforementioned mutations compared to the MpDCL1a wild-type sequence. For further analyses we focused on Mp*dcl1a-6*^*ge*^, as it exhibited severe developmental aberrations due to the genomic alteration that most likely affected the proper folding of MpDCL1a protein. The RES III domain of MpDCL1a (position 379–557) is predicted to be involved in endonucleolytic cleavage to generate double-stranded fragments and any change in the amino acid sequence affecting the secondary protein structure might impair the function of MpDCL1a and lead to faulty pri-miRNA processing.

Sequencing of mutated sites using plants from the G2 generation (Figure 2B) confirmed the generation of premature stop codons in Mp*dcl1b*^*ge*^, Mp*dcl3*^*ge*^, and Mp*dcl4*^*ge*^ lines (Supplementary Table 2). These lines together with Mp*dcl1a-6*^*ge*^ were further phenotypically examined. For this purpose, single gemmae of one male and one female line of Mp*dcl1b*^*ge*^, Mp*dcl3*^*ge*^, Mp*dcl4*^*ge*^ and WT control were plated on standard medium and observed for at least 21 days after gemmae germination (DAG). Only a female line of Mp*dcl1a-6*^*ge*^ was used in phenotypic analysis as we could not obtain a male line with the aforementioned mutations in the Mp*DCL1a* gene. While Mp*dcl1a-6*^*ge*^ and Mp*dcl3*^*ge*^ lines displayed severe developmental alterations compared to the WT, Mp*dcl1b*^*ge*^ and Mp*dcl4*^*ge*^ grew WT-like (Figure 2C). Mp*dcl1a-6*^*ge*^ showed an overall stunted, callus like growth under control conditions and did not develop gemmae cups but still produced gemmae on the dorsal side of the thallus (Figure 3A).

**Figure 3.**
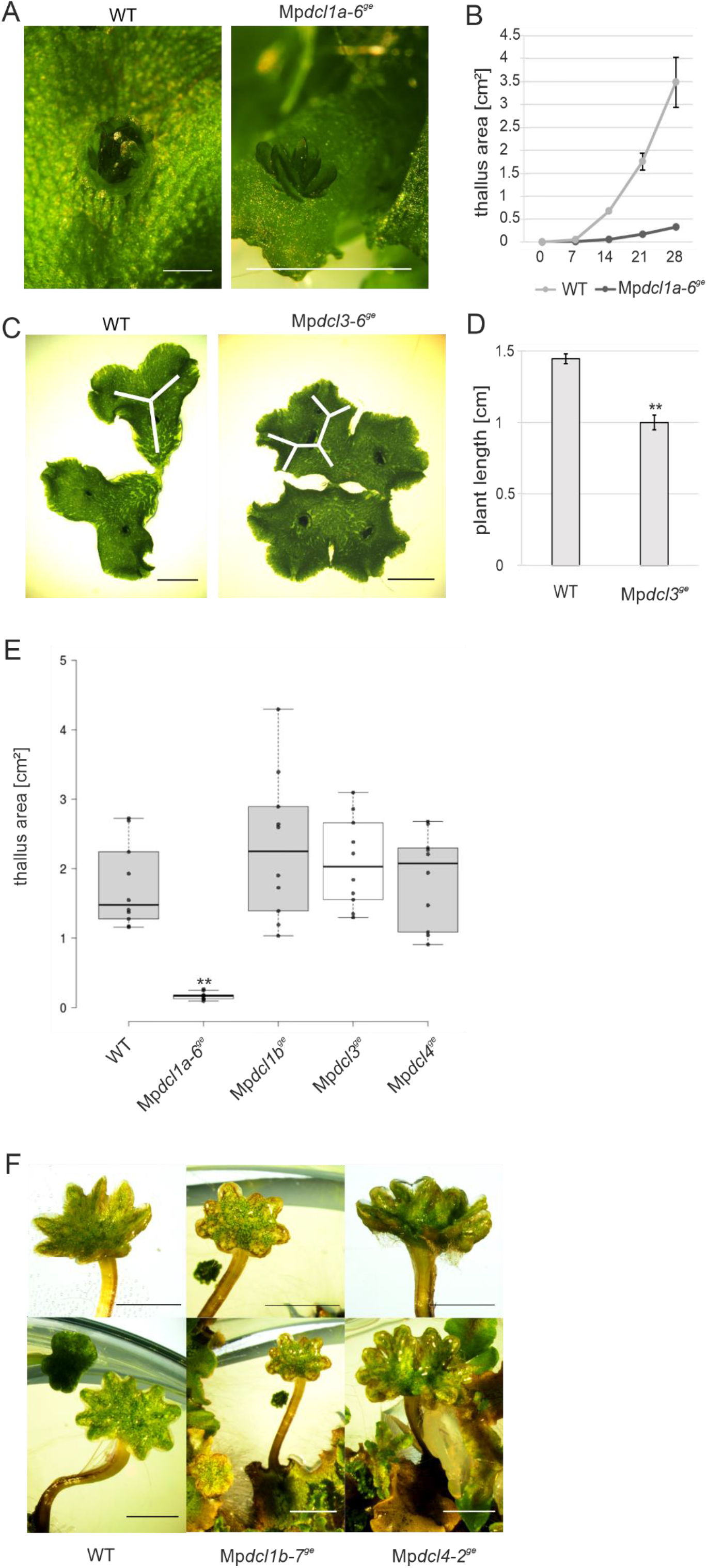
Phenotypic analysis of Mp*dcl*^*ge*^ mutant lines. **(A)** Close-up of a WT gemmae cup and gemmae from Mp*dcl1a-6*^*ge*^ (♀) that develop on the dorsal thallus side from 7 weeks old plants grown under control conditions on standard medium; scale bar 1 mm. **(B)** Thallus area of WT (♀/♂; 50/50) and Mp*dcl1a-6*^*ge*^ (♀) grown on standard medium under control conditions at 0, 7, 14, 21 and 28 days after gemmae germination (DAG), (n = 10, error bars indicate ± SEM). **(C)** Branching of WT and Mp*dcl3-6*^*ge*^, 21 DAG grown on standard medium under control conditions; scale bar 5 mm. **(D)** Total plant length (cm) of WT and Mp*dcl3*^*ge*^ (n = 10; ♀/♂; 50/50) 14 DAG. Statistically significant Student’s *t*-test (p < 0.001) is marked by two asterisks, error bars depict ± SEM. **(E)** Box plot of the thallus area of 21 DAG Mp*dcl1a-6*^*ge*^ (♀) as well as ♀ and ♂ (50/50) WT, Mp*dcl1b*^*ge*^, Mp*dcl3*^*ge*^, Mp*dcl4*^*ge*^ lines grown on standard medium under control conditions, n = 10. Statistically significant results of Student’s *t*-test between WT and mutant lines (p < 0.001) are marked with two asterisks. **(F)** Antheridiophores from WT, Mp*dcl1b-7*^*ge*^ and Mp*dcl4-2*^*ge*^ after 12 weeks growth in far-red light conditions (FRL). Upper panel: close-ups, lower panel: complete antheridiophore; scale bar 5 mm.

A closer look at the gemmae development in the Mp*dcl1a-6*^*ge*^ mutant revealed that they grow drastically slower than WT plants at 7 DAG and do not show a typical dichotomous thallus branching, but form a callus-like structure even after 21 DAG (Figure 2C, 3B). However, Mp*dcl3*^*ge*^ showed a slightly stunted thallus growth resulting in a reduced thallus length compared to WT. Interestingly, compared to the WT Mp*dcl3*^*ge*^ plants showed an increase in thallus branching events resulting in a distinct rosette shape at 21 DAG (Figure 3C). Even though this decrease in the thallus arm length of Mp*dcl3*^*ge*^ at 14 DAG was statistically significant (p < 0.001) (Figure 3D), this difference between Mp*dcl3*^*ge*^ (mean 2.09 cm^2^ ± SEM 0.2) and WT (mean 1.75 mm^2^ ± SEM 0.19) thallus area was no longer statistically significant (p = 0.243) at 21 DAG. Further statistical analysis of all respective mutant *DCL* lines at 21 DAG revealed that the thallus area of Mp*dcl1a-6*^*ge*^ was significantly reduced to an average of 0.17 cm^2^ (SEM ± 0.02) in comparison to WT (p < 0.001) (Figure 3E). Thallus growth of Mp*dcl1b*^*ge*^ (mean 2.3 cm^2^ ± SEM 0.33) and Mp*dcl4*^*ge*^ (mean 1.86 cm^2^ ± SEM 0.21) lines exhibited no significant changes compared to WT (mean 1.8 cm^2^ ± SEM 0.19) under control conditions (Figure 3E) (Mp*dcl1b*^*ge*^ p = 0.165; Mp*dcl4*^*ge*^ p = 0.723).

Consequently, these phenotypic analyses suggest an important function for Mp*DCL1a*, through sRNAs that are processed by MpDCL1a, in gemmae germination, thallus development and gemmae cup formation. Even though Mp*dcl3*^*ge*^ did not show any significant changes compared to WT at 21 DAG, the accelerated branching of Mp*dcl3*^*ge*^ in early developmental stages and different branching patterns suggest an essential role for Mp*DCL3* and associated sRNAs during thallus development. However, Mp*dcl1b*^*ge*^ and Mp*dcl4*^*ge*^ despite higher phenotypic variation, did not show any alterations in thallus development compared to WT. While Mp*DCL1b* and Mp*DCL4* are expressed at low levels in sporelings, gemmalings and adult thallus, they both are expressed at high levels in the antheridium (eFP browser, http://bar.utoronto.ca/efp_marchantia/cgi-bin/efpWeb.cgi) and mid-tier levels in the antheridiophores. Thereby, rather than developmental aberrations in sporelings or gemmalings, defects may be expected in antheridiophore development or antheridium formation due to mutations in Mp*dcl1b*^*ge*^ and Mp*dcl4*^*ge*^. Attempts to induce antheridiophores in Mp*dcl1b*^*ge*^ and Mp*dcl4*^*ge*^ together with WT plants by cultivation under far-red light supplemented light conditions resulted in smaller and deformed antheridiophores (Figure 3F). Surprisingly, no antheridiophores could be induced in Mp*dcl3*^*ge*^ and hence it can be suggested that MpDCL1b, MpDCL3 and MpDCL4, through associated sRNAs, play important roles in antheridiophore induction.

### Mutation in Mp*DCL1b* resulted in increased salt tolerance

In comparison to *P. patens, M. polymorpha* shows a lower salt tolerance and its survival rate drops drastically when grown on solid media supplemented with 50 mM NaCl (Frank *et al*., 2005, Tanaka *et al*., 2018). We recently showed that the abolishment of the *PpDCL1a*/miR1047 autoregulatory feedback loop leads to salt hypersensitivity in *P. patens* indicating the importance of DCLs and sRNAs in salt stress tolerance (Willmann and Poethig, 2007, Barciszewska-Pacak *et al*., 2015, Arif *et al*., 2022). Previous reports also revealed that sRNAs are crucial for abiotic stress tolerance mechanisms in various plant species (Fu *et al*., 2017, Habermann *et al*., 2020, Tiwari *et al*., 2020, Tiwari *et al*., 2021) and due to the anticipated adjustment and/or misregulation of sRNA biogenesis and sRNA-directed gene regulatory networks, we hypothesized that plants with mutated Mp*DCLs* would have altered abiotic stress tolerance. High salinity is one of the most severe abiotic stresses that induces global transcriptomic changes and adversely affects plant growth and development (Golldack *et al*., 2011, Fu *et al*., 2017, Tanaka *et al*., 2018). For example, transcription factors of the MYB, bHLH, and bZIP families are differentially expressed upon salt treatment in *A. thaliana* (Golldack *et al*., 2011) and *M. polymorpha* (Tanaka *et al*., 2018).

To test whether mutations in Mp*DCL* genes lead to altered salt stress response, we plated multiple gemmae (Ø 118.9 gemmae per plate) from Mp*dcl1a-6*^*ge*^, male and female Mp*dcl1b*^*ge*^, Mp*dcl3*^*ge*^, and Mp*dcl4*^*ge*^ lines as well as male and female WT lines, on standard medium supplemented with 250 mM NaCl and grew them for 14 d under standard conditions (Figure 4A). After 14 d under salt stress conditions, we counted all plants showing growth and chlorophyll production on each plate and calculated the survival rate of the respective lines. At 14 DAG male and female WT lines showed an average survival rate of 30.03 % (Figure 4B) which is in line with a previous study (Tanaka *et al*., 2018). However, an overall decrease in survival rates to 20.84 % and 14.30 % was observed for Mp*dcl3*^*ge*^ and Mp*dcl4*^*ge*^ lines, respectively (Figure 4B). Despite the observed decrease in both lines, only the decrease in Mp*dcl4*^*ge*^ lines upon salt treatment was statistically significant (p = 0.004). On the other hand, both Mp*dcl1a-6*^*ge*^ *and* Mp*dcl1b*^*ge*^ displayed an enhanced salt stress tolerance when compared to WT the other mutant lines and showed an average survival rate of 45.03 % (p = 0.003) and 68.90 % (p < 0.001), respectively (Figure 4B). To sum up, mutations in Mp*DCL1a*, Mp*DCL1b* and Mp*DCL4* resulted in altered survival rates upon salt treatment. While MpDCL4 and its associated sRNAs positively regulate salt stress adaptation in *M. polymorpha*, MpDCL1a, MpDCL1b and those sRNAs that are controlled by these proteins are most likely negative regulators of salt stress adaptation. In particular this might explain why *M. polymorpha* wild-type which harbors functional MpDCL1a, MpDCL1b and MpDCL4 does not tolerate high levels of salt stress and further investigations are required to understand how these mechanisms evolved during the water-to-land transition.

**Figure 4.**
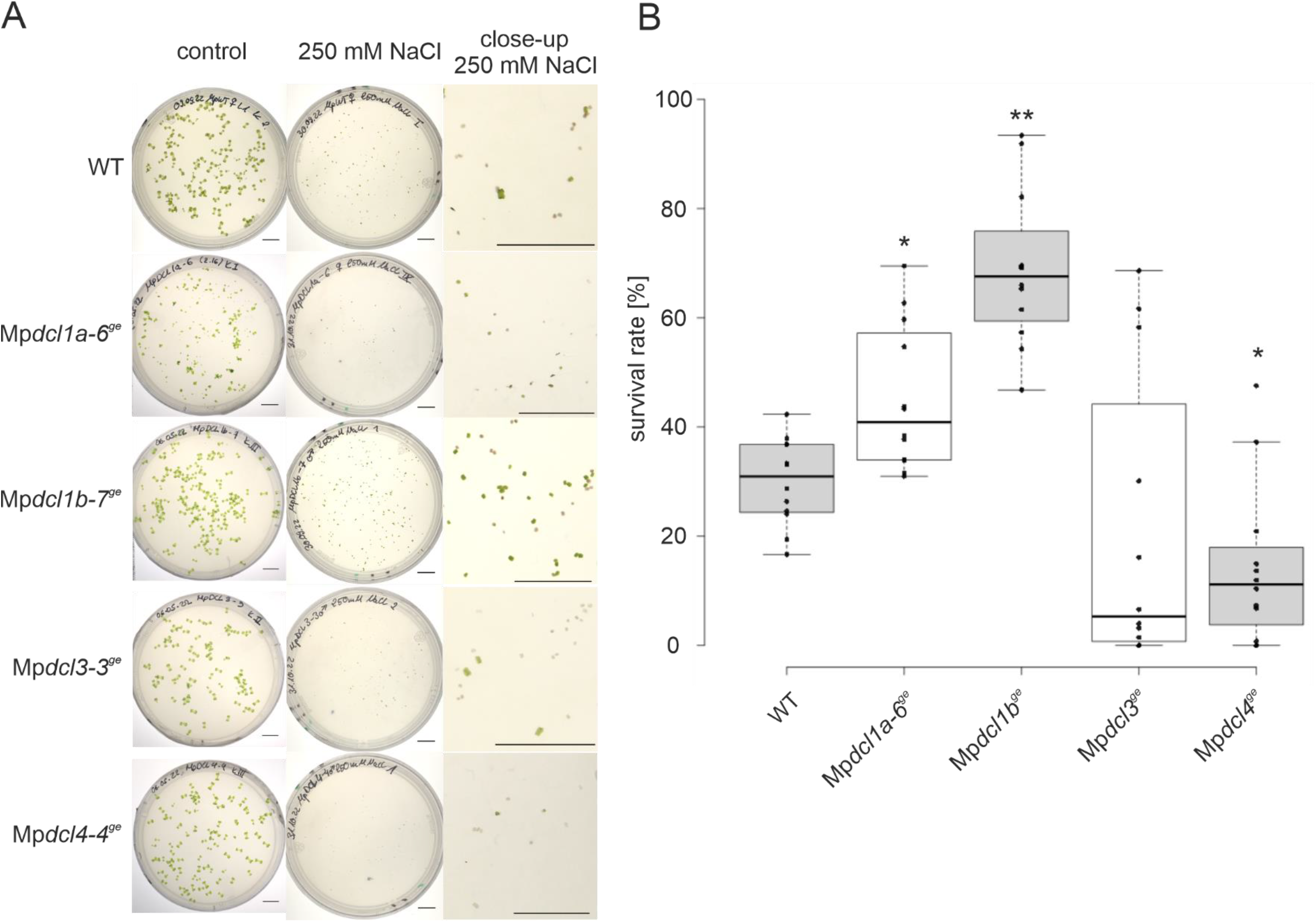
Growth of WT and Mp*dcl*^*ge*^ mutant lines on high salt concentration. **(A)** WT, Mp*dcl1a-6*^*ge*^, Mp*dcl1b-7*^*ge*^, Mp*dcl3-3*^*ge*^, and Mp*dcl4-4*^*ge*^ grown for 14 days after gemmae germination (DAG) on standard medium (left) and medium supplemented with 250 mM NaCl (middle). The right column depicts close-ups of plates with the NaCl-supplemented medium; scale bar 1 cm. **(B)** Box plot of survival rates of Mp*dcl1a-6*^*ge*^ (♀), as well as ♀ and ♂ (50/50) WT, Mp*dcl1b*^*ge*^, Mp*dcl3*^*ge*^, Mp*dcl4*^*ge*^ lines on medium containing 250 mM NaCl 14 DAG. Statistically significant results of Student’s *t*-test between WT and the respective Mp*dcl*^*ge*^ mutant lines with p < 0.001 are marked with two asterisks and p < 0.05 with one asterisk.

### Mp*dcl1a-6*^*ge*^, Mp*dcl3*^*ge*^ and Mp*dcl4*^*ge*^ lines display ABA hypersensitivity

Numerous miRNAs are differentially regulated in response to phytohormone treatment and upon abiotic stress (Khraiwesh *et al*., 2010, Khraiwesh *et al*., 2012). Abscisic acid (ABA) is the central mediator of stress signaling responses (Hauser *et al*., 2011, Arif *et al*., 2018), hence alterations in the various sRNA biogenesis pathways due to mutations in Mp*DCL* genes might affect ABA signaling as well as the sensitivity of the mutant lines to ABA treatment. We analyzed whether the response to ABA in the Mp*dcl1a-6*^*ge*^, Mp*dcl1b-6*^*ge*^, Mp*dcl1b-7*^*ge*^, Mp*dcl3-3*^*ge*^, Mp*dcl3-6*^*ge*^, Mp*dcl4-1*^*ge*^ and Mp*dcl4-4*^*ge*^ lines would be affected by the loss of the respective functional DCL protein and the subsequent changes in the sRNA and mRNA repertoire upon exogenous ABA treatments. We inoculated single gemmae (n = 10) of the respective Mp*DCL* mutant lines (both male and female lines, with the exception of Mp*dcl1a*^*ge*^, together with WT) on standard medium supplemented with 10 μM ABA. Gemmae were grown under standard conditions for at least 21 DAG before thallus area was measured and compared with WT controls. In response to ABA, WT showed reduced growth at 21 DAG (mean 0.39 cm^2^; SEM ± 0.02) and a 5.5-fold decrease in thallus area compared to the untreated WT (Figure 5A). These observations are in accordance with a previous study where ABA treatment led to a delay in gemmae germination and stunted growth (Eklund *et al*., 2018). Even though Mp*dcl1a-6*^*ge*^ displayed stunted growth under normal conditions, its growth (i.e., thallus area) decreased by 8.6-fold in response to ABA. This decrease in biomass in comparison to ABA-treated WT is statistically significant (p < 0.001) (Figure 5A, 5B). While Mp*dcl1b*^*ge*^ mutant lines showed reduced growth compared to untreated Mp*dcl1b*^*ge*^ lines, this ABA sensitivity was not statistically significant (p = 0.088) most likely due to observed high variability between plants (Figure 5B). Moreover, Mp*dcl3*^*ge*^ and Mp*dcl4*^*ge*^ lines showed ABA hypersensitivity like Mp*dcl1a-6*^*ge*^: 8.6-fold and 6.6-fold decreases in thallus area were recorded in Mp*dcl3*^*ge*^ (p < 0.001) and Mp*dcl4*^*ge*^ (p = 0.008), respectively (Figure 5B). Consequently, it can be proposed that all MpDCLs except MpDCL1b and those sRNAs associated with these proteins are involved in ABA signaling and stress response in *M. polymorpha*.

**Figure 5.**
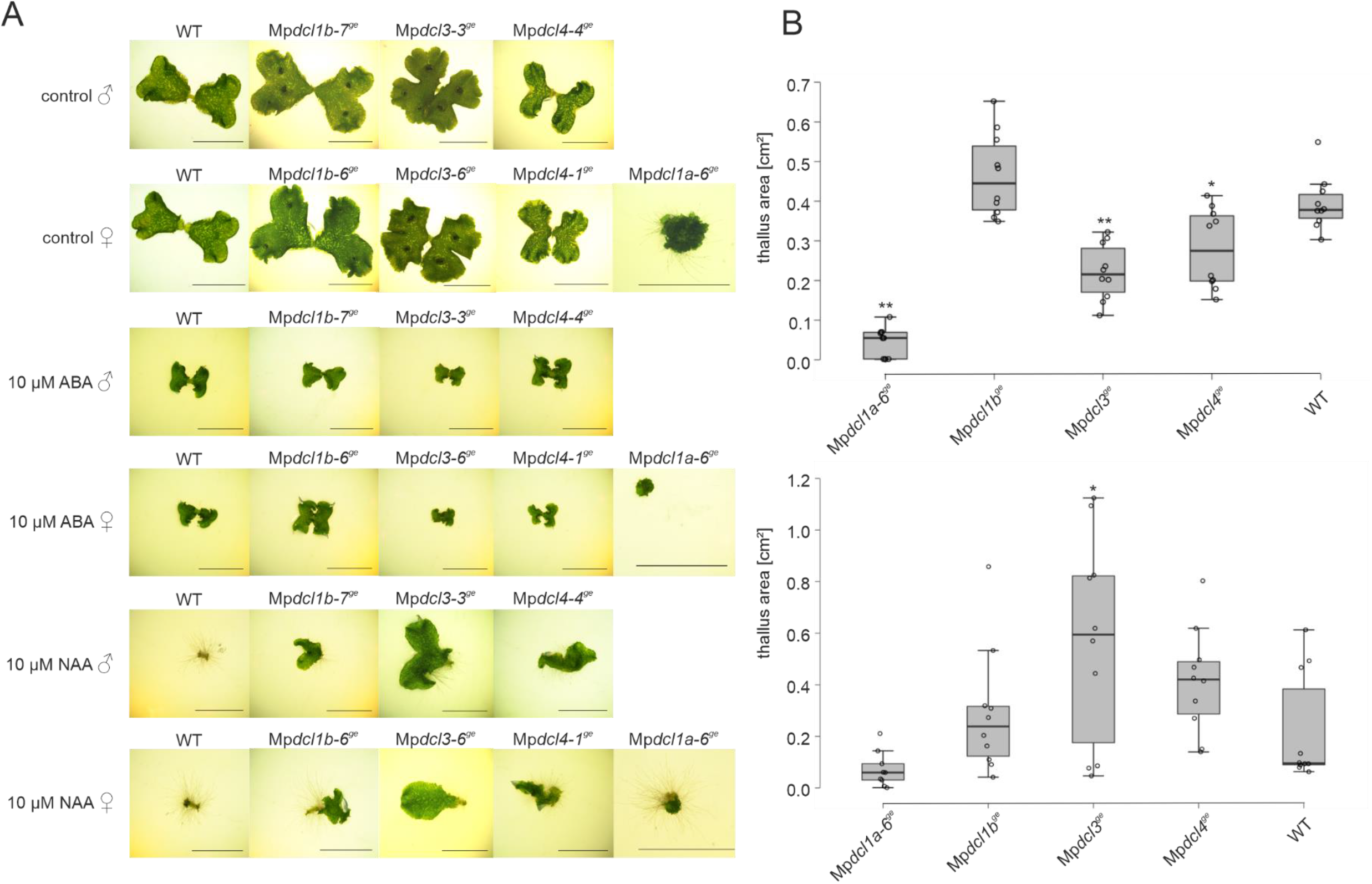
Phytohormone treatment of mutant Mp*dcl*^*ge*^ lines. **(A)** Male and female lines of WT and Mp*dcl1b*^*ge*^, Mp*dcl3*^*ge*^, and Mp*dcl4*^*ge*^ lines as well as the Mp*dcl1a-6*^*ge*^ female line grown for 21 days after gemmae germination (DAG) on standard medium and medium supplemented with 10 μM ABA or 10 μM NAA; scale bar 1 cm. **(B)** Box plots of thallus area (cm^2^) of male and female (50/50) WT, Mp*dcl1b*^*ge*^, Mp*dcl3*^*ge*^, and Mp*dcl4*^*ge*^ lines grown on standard medium supplemented with 10 μM ABA (upper panel) and 10 μM NAA (lower panel) for 21 DAG under control conditions; n=10 (5 ♀ and 5 ♂ per line) except for Mp*dcl1a-6*^*ge*^ (n = 9 all ♀). Statistically significant results of Student’s *t*-test between WT and the respective mutant line are marked with two asterisks (p < 0.001) and or one asterisk (p < 0.05).

### Mp*dcl3*^*ge*^ phenotype is rescued by auxin treatment

Auxin regulates plant growth and differentiation through coordinated changes in gene expression and assists the transition from 2D to 3D growth in *M. polymorpha* (Flores-Sandoval *et al*., 2015). In response to exogenous auxin, while thallus growth is inhibited, rhizoidal initiation and elongation are observed in the liverwort *M. polymorpha* (Ishizaki *et al*., 2012, Kato *et al*., 2017). Similar changes can be observed in various plant species upon auxin treatment. For instance, auxin induces chloronema to caulonema transition, inhibits bud formation and controls rhizoid and gametophore development in *P. patens* (Jang and Dolan, 2011, Thelander *et al*., 2018). Moreover, miRNAs have been shown to modulate auxin signaling and thereby plant development and stress responses (Navarro *et al*., 2006, Jodder, 2020). To investigate whether mutations in Mp*DCLs* and an altered sRNA repertoire result in impaired auxin response and developmental aberrations, we treated mutant as well as wild-type plants with exogenous NAA. Ten gemmae were grown on standard medium supplemented with 10 μM NAA and the visible thallus area was measured 21 DAG. In wild-type exogenous NAA treatment induced enhanced rhizoid formation and inhibited thallus growth significantly (p < 0.001, in comparison with untreated WT) (Figure 5A), which is in line with previous findings in *M. polymorpha* (Ishizaki *et al*., 2012, Flores-Sandoval *et al*., 2015, Kato *et al*., 2017). Upon exogenous NAA treatment, Mp*dcl1a-6*^*ge*^ plants grew significantly slower (p = 0.002) compared to untreated Mp*dcl1a-6*^*ge*^ plants (21 DAG). In addition, Mp*dcl1a-6*^*ge*^ showed increased rhizoid production like auxin-treated wild-type plants (Figure 5A). However, the reduction in thallus area in Mp*dcl1a-6*^*ge*^ plants after NAA treatment was not statistically significant (p = 0.057) when compared to WT plants (Figure 5B). Treatment of Mp*dcl3*^*ge*^ with NAA led to overall growth inhibition (p < 0.001) that was significantly different from WT (p = 0.027). Rather than induced rhizoid formation at the expense of inhibited thallus growth, Mp*dcl3*^*ge*^ plants resembled the WT under control conditions indicating a NAA insensitive phenotype for these mutants (Figure 5A). Reduced thallus growth (p < 0.001) and increased rhizoid formation were also observed in Mp*dcl1b*^*ge*^ and Mp*dcl4*^*ge*^ upon exogenous NAA treatment (Figure 5B), but they did not exhibit NAA hypo- or hypersensitivity as reductions in thallus growth in these mutant lines were not statistically significant compared to the growth reduction in WT (Mp*dcl1b*^*ge*^ p = 0.517; Mp*dcl4*^*ge*^ p = 0.056). In conclusion, due to the NAA insensitivity observed in the Mp*dcl3*^*ge*^ mutant, we propose that MpDCL3 and its associated sRNAs are involved in auxin signaling, essential for *M. polymorpha* development.

## Discussion

DCL proteins are functionally conserved proteins that play crucial roles during the water-to-land transition. The adaptation to challenging environments increased and diversified DCLs in land plants (Wang *et al*., 2021). While DCL1a, DCL2, DCL3 and DCL4 are major members that are present in the majority of land plants, some monocots also encode DCL5 (Wang *et al*., 2021). In line with previous studies, our phylogenetic analysis confirmed that *M. polymorpha* harbors four DCLs: MpDCL1a, MpDCL1b, MpDCL3 and MpDCL4. Except MpDCL1b, all other three DCLs contain RESIII, Helicase C, PAZ, Dicer-dimer, two adjacent RNase III, and at least one double-strand RNA binding domain. These three DCLs are highly similar to homologs in *P. patens*. While MpDCL1a and MpDCL3 have the same domain structure as their counterparts PpDCL1a and PpDCL3 in *P. patens*, MpDCL4 contains an additional DND1_DSRM domain compared to PpDCL4. Interestingly, MpDCL1b harbors RESIII, PAZ and two adjacent RNase III domains, whereas PpDCL1b contains additional Helicase C, Dicer-dimer, and DND1_DSRM domains. The absence of functionally important domains in MpDCL1b raises the question whether it is a pseudogene. Our comparative phylogenetic analysis revealed that *A. agrestis* and *S. cucullata* encode DCLs that lack important domains like MpDCL1b. Hence, we propose that MpDCL1b and PpDCL1b evolved independently from each other and MpDCL1b might fulfill different roles in sRNA biogenesis.

While DCL1 homologs such as PpDCL1a and AtDCL1 are essential for miRNA biogenesis, the generation of *DCL1* deletion mutants results in either developmentally arrested plants like in *P. patens* (Khraiwesh *et al*., 2010) and *O. sativa* (Liu *et al*., 2005) or embryonic lethality like in *A. thaliana* (Henderson *et al*., 2006, Arif *et al*., 2013). Since we could not identify any Mp*dcl1a*^*ge*^ mutant line with large genomic rearrangements or early stop codons, we propose potential lethality of Mp*DCL1a* null mutants. The detected mutations in the Mp*DCL1a* gene that led to specific amino acid exchanges resulted in severe developmental abnormalities, including stunted growth and inhibition of gemmae cup formation similar to the drastic phenotypic abnormalities in the Δ*PpDCL1a* mutant line (Khraiwesh *et al*., 2010, Arif *et al*., 2022). Our comprehensive phylogenetic analysis as well as observed phenotypic aberrations in the mutant lines suggest that MpDCL1a is a PpDCL1a homolog and most likely modulates miRNA biogenesis in *M. polymorpha*. Further investigations including mRNA and sRNA sequencing are required to reveal the role of MpDCL1a in sRNA biogenesis. However, due to the mutations in regions encoding the restriction enzyme subunit (RESIII domain) within MpDCL1a, it can be speculated that cleavage might be error-prone and inefficient during miRNA biogenesis.

Previous studies already showed the impact of specific miRNAs on developmental programs in *M. polymorpha*. For instance, ectopic expression of *MIR166* in *M. polymorpha* led to stunted growth with ventral curling of the thallus branches and shorter gemmae cups (Tsuzuki *et al*., 2016) whereas overexpression of *MIR319* resulted in the complete loss of gemmae cups, but normal growth of gemmae (Tsuzuki *et al*., 2016, Pietrykowska *et al*., 2022).

Mutations in the Mp*DCL1b* gene neither affected gemmae germination nor thallus development. However, Mp*dcl1b*^*ge*^ lines showed an increased salt-stress tolerance compared to the WT and the other *MpDCL* mutant lines. Interestingly, the expression data taken from the eFP browser expression atlas (http://bar.utoronto.ca/efp_marchantia/cgi-bin/efpWeb.cgi) show a slight downregulation of Mp*DCL1b* upon salt treatment (Tan *et al*., 2022). Even though these lines show an ABA-hyposensitive phenotype, statistical analysis failed to show any significance due to high variation in the observed lines. On the other hand, *Mpdcl4*^*ge*^ lines displayed hypersensitivity to salt most likely due to altered sRNA-dependent regulation of salt-responsive genes (Kasschau *et al*., 2007).

DCL3 is responsible for the generation of short interfering RNAs (siRNAs) in land plants and these siRNAs participate in transcriptional silencing via regulation of epigenetic modifications like RNA-dependent DNA methylation or histone modifications (Henderson *et al*., 2006, Moura *et al*., 2019). These siRNAs are derived from repetitive regions and transposable elements. In *P. patens*, PpDCL3 is required for the accumulation of 22–24 nt siRNAs and Δ*PpDCL3* gene deletion mutants display an accelerated gametophore development (Cho *et al*., 2008). In line with this role of PpDCL3 in *P. patens*, Mp*dcl3*^*ge*^ lines developed shorter thallus branches and showed accelerated branching compared to the WT. Moreover, the aberrant Mp*dcl3*^*ge*^ phenotype was reverted to a wild-type-like growth by exogenous auxin application suggesting a potential role of MpDCL3 and its associated sRNAs in the modulation of auxin-responsive genes. Auxin acts in the establishment of the dorsoventral polarity, rhizoid and gemmae cup formation, developmental stage transition and determines the overall body plan as well as branching patterns in *M. polymorpha* (Flores-Sandoval *et al*., 2015, Flores-Sandoval *et al*., 2018b). Flores-Sandoval et al. (2015) reported an increased branching rate by overexpressing *AUXIN RESPONSE FACTOR 1* (Mp*ARF1*) and repressing *TOPLESS* (Mp*TPL*) (Flores-Sandoval *et al*., 2015). Moreover, the control of Mp*ARFs* is mediated by miRNAs since miR160 regulates C class *ARF* transcripts in *M. polymorpha* (Flores-Sandoval *et al*., 2018a). It has been shown that Mp*arf3* mutants display an inhibition of differentiation and developmental transitions in their life cycle and Mp*miR160* mutants do not produce gametangiophores upon far-red light supplementation (Flores-Sandoval *et al*., 2018a). Thus, we propose that MpDCL and its associated sRNA most likely regulate auxin-responsive genes including Mp*ARFs* and Mp*TPLs* that is also supported by the lack of antheridiophore development in Mp*dcl3*^*ge*^ mutant lines.

Phenotypic analyses of Mp*dcl4*^*ge*^ mutant lines under control conditions and upon exogenous phytohormone treatment did not reveal differences compared to the WT. These observations differ widely from *P. patens* where loss of the *PpDCL4* gene resulted in severe developmental defects, gravitropism insensitivity and sterility (Khraiwesh *et al*., 2010). A detailed analysis using the available eFP expression atlas (http://bar.utoronto.ca/efp_marchantia/cgi-bin/efpWeb.cgi) revealed that Mp*DCL4* is barely expressed during thallus development while it is highly expressed in gametangiophores, especially in the spermatid mother cells and the antheridial cavity. This tissue-specific expression hints at a possible function of Mp*DCL4* and its associated sRNAs during spermatozoa formation and a possible sterility of Mp*dcl4*^*ge*^ lines that needs to be confirmed. Nevertheless, our attempts to induce antheridiophores in Mp*dcl4*^*ge*^ resulted in production of smaller, less and malformed antheridiophores compared to WT. The discrepancy with respect to phenotypic aberrations between Mp*dcl4*^*ge*^ and Δ*PpDCL4* lines might be due to the role and importance of MpDCL4 in trans-acting small interfering RNA (ta-siRNA) biogenesis in *M. polymorpha*. Three miRNAs responsible for ta-siRNA biogenesis as well as four *TAS* gene families are present in *A. thaliana* (Allen *et al*., 2005, Arif *et al*., 2012) while all ta-siRNAs originate from four *TAS3* precursors in *P. patens* (Axtell *et al*., 2006, Arif *et al*., 2012). In contrast, only one *TAS* locus (Mp*TAS3*) with two miR390 binding sites has been identified in *M. polymorpha* (Tsuzuki *et al*., 2016). In the same study the authors detected 20–21 nt siRNA in antheridiophores derived from two additional loci without complementary miRNAs triggering their expression. These ta-siRNAs are designated as sex-specific phasing siRNAs (SS-phasiRNAs) and their processing is proposed to be executed by MpDCL4 (Tsuzuki *et al*., 2016). The lack of detectable ta-siRNAs together with the lack of phenotypic alterations under normal growth conditions in Mp*dcl4*^*ge*^ mutants suggest that ta-siRNAs are not required to control *M. polymorpha* development. On the other hand, ta-siRNAs regulate bud formation and transition from 2D to 3D growth in *P. patens* (Cho *et al*., 2012). Surprisingly, we detected decreased salt-stress tolerance and increased ABA sensitivity in Mp*dcl4*^*ge*^ mutant lines. We speculate that MpDCL4 is responsible for processing of phasing siRNAs (phasiRNAs) and ta-siRNAs and a change in the biogenesis of these sRNAs may influence survival at high salt concentrations and phytohormone sensitivity. Hence, we propose that MpDCL4 might fulfill similar functions as PpDCL4, but the role and importance of ta-siRNAs can differ between the moss *P. patens* and the liverwort *M. polymorpha*.

## Conclusion

Elucidation of DCL functions via comparative analysis can help us to understand the evolutionary origin of DCL proteins in land plants. In this study we focused on the role of the four encoded DCL proteins MpDCL1a, MpDCL1b, MpDCL3 and MpDCL4 in *M. polymorpha*. While we observed phenotypic similarities between *M. polymorpha* and *P. patens DCL* mutant lines, distinct phenotypic differences were also recorded especially in Mp*dcl3*^*ge*^ and Mp*dcl4*^*ge*^ mutant lines. Moreover, we found that MpDCL1b and its associated sRNAs act as negative regulators of salt tolerance, whereas MpDCL4 and its associated sRNAs act as a positive regulators of salt stress adaptation. Consequently, we conclude that MpDCLs are important for proper plant growth and also have specialized functions that remain to be elucidated in detail in the future.

## Material and Methods

### Phylogenetic tree of DICER-LIKE proteins

To obtain homologous protein sequences of MpDCL1a (Mp7g12090), MpDCL1b (Mp6g09830), MpDCL3 (Mp1g02840) and MpDCL4 (Mp7g11720) we used their sequences as queries to search in the databases Phytozome v13 (phytozome-next.jgi.doe.gov), MarpolBase (marchantia.info), Ensembl Plants re55 (plants.ensembl.org) as well as UniProt (re 2022_04; uniport.org) with BLASTp for reciprocal matches in the species *Anthoceros agrestis, Arabidopsis thaliana, Azolla filiculoides, Chara braunii, Chlamydomonas reinhardtii, Klebsormidium nitens, Medicago truncatula, Oryza sativa japonica, Physcomitrium patens, Salvinia cucullata, Selaginella moellendorfii, Spirogloea muscicola*, and *Zea mays*. All protein sequences (for accession numbers see Supplementary Table 1) were aligned using CLC Genomics Workbench v20.0.4 (Qiagen) and the phylogenetic tree was constructed with the neighbor-joining method using 1000 bootstrap replicates. Only nodes with bootstrap values >50% are shown and nodes >90% are highlighted.

### Cultivation of *Marchantia polymorpha*

We have used *M. polymorpha ssp. ruderalis* ecotype BoGa obtained from the Botanical Garden of the University of Osnabrück (Germany) throughout our studies. All plants were grown under sterile conditions on standard solid half-strength Gamborg B5 medium including vitamins (Duchefa) (pH 5.5, 1.4 % Agar-Agar, Kobe I) under long-day conditions (16L:8D) at 23°C and a light intensity of 85–100 μmol/m^2^s according to Althoff et al. (2014). Gametangiophore induction was performed according to Althoff et al. (2014) at 23°C.

### Generation of *M. polymorpha* mutant lines

*M. polymorpha* mutant lines were generated following the double CRISPR/Cas 9 approach described by Sugano et al. (2018). Double-strand breaks were induced resulting in sequence deletions or nucleotide insertions leading to frameshift mutations of the targeted gene loci Mp*DCL1a*, Mp*DCL1b*, and Mp*DCL3* and Mp*DCL4* (Supplementary Table 2). All synthetic gRNAs were designed with CRISPOR (http://crispor.org) (Concordet and Haeussler, 2018) and are targeting the first two exons of the Mp*DCL* locus or regions that code for functional domains. Cloning was carried out as described in Sugano et al. (2018) using the vectors pMpGE_EN03, pMpGE010, and pMpGE011 each encoding different gRNAs (for gRNAs see Supplementary Table 2 and 3) and used in combination with each other (Sugano *et al*., 2018). *A. tumefaciens* strain C58C1 pGV2260 mediated transformation was performed as previously described (Ishizaki *et al*., 2008) with minor modifications. 7 d old WT sporelings were co-cultivated for 3 d with two *A. tumefaciens* lines, one carrying the first gRNA in pMpGE010 and the other carrying the second gRNA in pMpGE011, performing all transformations with two gRNAs for the same Mp*DCL* in parallel. The transformed plants were plated on cellophane covered standard medium supplemented with 0.5 μM chlorosulfuron (Sigma), 10 mg/L hygromycin (Hygromycin B-solution; Roth), and 100 mg/L cefotaxime (Cefotaxime sodium; Duchefa) for two weeks before transferring the transformants to standard medium supplemented with 1 % glucose (Roth) and cultivating them for two weeks. Afterward, a second round of selection was performed for three weeks before further analysis. All transformed T^1^ plants were screened by PCR (for oligonucleotides see Supplementary Table 3) and isogenic lines were obtained by propagating single gemmae to avoid potential chimeric lines. Regions that were targeted by the respective gRNAs were further sequenced to identify mutations and/or frameshifts caused by the CRISPR/Cas9 approach. With the exception of Mp*DCL1a* (only female), one female and one male mutant line was chosen per Mp*DCL* gene and the lines Mp*dcl1a-6*^*ge*^, Mp*dcl1b-6*^*ge*^, Mp*dcl1b-7*^*ge*^, Mp*dcl3-6*^*ge*^, Mp*dcl3-3*^*ge*^, Mp*dcl4-1*^*ge*,^ and Mp*dcl4-4*^*ge*^ were used for all further analyses presented (Supplementary Table 2).

### DNA and RNA extraction

DNA extractions were performed with 0.5–1 cm^2^ thallus pieces and 200 μl of extraction buffer (100 mM Tris-HCl, 1 M KCl, 10 mM EDTA, pH 9.5). The tissue was disrupted with a Qiagen TissueLyser II and DNA was extracted according to Edwards et al. (1991). RNA extraction was performed with TRIzol™ Reagent (Invitrogen) following the manufacturer’s protocol out of 50–100 mg plant material. RNA samples were dissolved in nuclease-free water and stored at −80°C. RNA concentrations and quality were determined via NanoDrop™ 2000/2000c spectrophotometer (Invitrogen) and RNA integrity was confirmed by agarose gel electrophoresis.

### Phenotypic analyses

For phenotypic analyses single gemmae were placed on standard solid medium (control conditions) or medium supplemented with 10 μM 1-naphthylacetic acid (NAA) (1mg/ml, Sigma-Aldrich) or 10 μM 2-*cis*,4-*trans*-Abscisic acid (ABA) (Sigma-Aldrich), respectively. Images of at least five plants per mutant line were taken with a SMZ 1500 stereomicroscope and a DS-U3 camera (Nikon) at 0, 7, 14, 21 days after gemmae germination. Images of complete plates were taken with the EOS 800D camera (Canon). Area measurements for all individual plants were performed using ImageJ (Schneider *et al*., 2012) with the area measurement tool set to limit to threshold. Area was measured after setting the threshold covering the visible thallus area of the respective plant. For statistical analysis, average values of the measured area from at least two different mutant lines of the targeted genes Mp*DCL1a, 1b, 3* and *4* were compared with at least two independent WT lines with a sample size (n) of 10. For comparison of the thallus arm length of WT and Mp*dcl3*^*ge*^ complete plant length at 7 DAG, starting and ending at the first branching point of either side, was measured with ImageJ. Measured thallus area values and thallus length of the respective Mp*dcl*^*ge*^ mutant lines were tested for normal distribution with the Shapiro-Wilk test and statistical analysis of overall growth was performed using Student’s *t*-test when samples were normally distributed and Welch *t*-test when the samples were not normally distributed. The statistical analysis was performed with the statistical program JASP (JASP 0.9.0.1). Box plots were generated with JASP and the web-tool BoxPlotR (http://shiny.chemgrid.org/boxplotr/) (Spitzer *et al*., 2014).

### Salt stress tolerance assay

To analyze the salt tolerance, gemmae of one male and one female WT line as well as gemmae from all mutant lines (Mp*dcl1a-6*^*ge*^, Mp*dcl1a-8*^*ge*^, Mp*dcl1b-6*^*ge*^, Mp*dcl1b-7*^*ge*^, Mp*dcl3-6*^*ge*^, Mp*dcl3-3*^*ge*^, Mp*dcl4-1*^*ge*^, and Mp*dcl4-4*^*ge*^) were plated on standard solid medium supplemented with 250 mM NaCl (average of 118.9 gemmae/plate, SEM ± 5.6, n = 6) for 14 d. Images of the plates were taken at 0 days and after 14 d. Surviving gemmae were counted with ImageJ (Schneider *et al*., 2012). A statistical comparison of the survival rate of male and female mutant lines and the WT lines was performed with Student’s *t*-test. When no significant differences between male and female lines could be detected, survival rates of male and female lines were pooled (n = 12) and statistical analysis of the survival rate of mutant lines compared to the WT was performed with Student’s *t*-test and by inequality of variance with Welch *t*-test with the statistical program JASP (JASP 0.9.0.1).

## Acknowledgements

This project was carried out in the framework of MAdLand (https://madland.science/, DFG Priority Program 2237). WF and EC are grateful for funding by the Deutsche Forschungsgemeinschaft (DFG; FR 1677/5-1).

## Author contributions

EC, OT and WF designed the research. EC, AO, and AG performed the research and NG supported *M. polymorpha* experiments. All authors analyzed the data. EC, OT and WT wrote the manuscript with input from SZ and NG. All authors reviewed the manuscript.

## Data availability statement

All data generated or analyzed during this study are included in this article and its supplementary information.

## SUPPLEMENTARY TABLES

***Supplementary Table 1***. *Sources of all protein sequences used to generate the phylogenetic tree by neighbor joining and their identified functional domains*

***Supplementary Table 2***. *Accession numbers of the targeted* MpDCLs, *gRNA sequences and description of the respective mutation*

***Supplementary Table 3***. *Oligonucleotides used in this study*

